# Regulation of olfactory-based sex behaviors in the silkworm by genes in the sex-determination cascade

**DOI:** 10.1101/2020.01.24.917906

**Authors:** Jun Xu, Wei Liu, Dehong Yang, Shuqing Chen, Kai Chen, Zulian Liu, Xu Yang, Jing Meng, Guanheng Zhu, Shuanglin Dong, Yong Zhang, Shuai Zhan, Guirong Wang, Yongping Huang

## Abstract

Insect courtship and mating depend on integration of olfactory, visual, and tactile cues. Compared to other insects, *Bombyx mori*, the domesticated silkworm, has relatively simple sexual behaviors as it cannot fly. Here by using CRISPR/Cas and electrophysiological techniques we found that courtship and mating behaviors are regulated in male silk moths by mutating genes in the sex determination cascade belonging to two conserved pathways. Loss of *Bmdsx* gene expression significantly reduced the peripheral perception of the major pheromone component bombykol by reducing expression of the product of the *BmOR1* gene which completely blocked courtship in adult males. Interestingly, we found that mating behavior was regulated independently by another sexual differentiation gene, *Bmfru.* Loss of *Bmfru* completely blocked mating, but males displayed normal courtship behavior. Lack of *Bmfru* expression significantly reduced the perception of the minor pheromone component bombykal due to the down regulation of *BmOR3* expression; further, functional analysis revealed that loss of the product of *BmOR3* played a key role in terminating male mating behavior. Our results suggest that *Bmdsx* and *Bmfru* are at the base of the two primary pathways that regulate olfactory-based sexual behavior.

**Author Summary:** The fundamental insect sexual behaviors, courtship and mating, result from successful integration of olfactory, vision, tactile and other complex innate behaviors. In the widely used insect model, *Drosophila melanogaster*, the sex determination cascade genes *fruitless* and *doublesex* are involved in the regulation of courtship and mating behaviors; however, little is known about the function of these sexual differentiation genes in regulating sex behaviors of Lepidoptera. Here we combine genetics and electrophysiology to investigate regulation pathway of sexual behaviors in the model lepidopteran insect, the domesticated silk moth, *Bombyx mori*. Our results support the presence of two genetic pathways in *B. mori*, named *Bmdsx*-*BmOR1*-bombykol and *Bmfru*-*BmOR3*-bombykal, which control distinct aspects of male sexual behavior that are modulated by olfaction. This is the first comprehensive report about the role of sex differentiation genes in the male sexual behavior in the silk moth.

## Introduction

Sex determination pathways control the sexually dimorphic traits of males and females, including sexual differentiation and sexual behavior [1]. The genetic cascades of primary signalling that underlie sex determination in insects have high diversity among species. In the model insect *Drosophila melanogaster*, sex determination is controlled hierarchically by X:A, *Sex-lethal* (*Sxl*), *transformer* or *transformer* 2 (*tra*/*tra2*), *doublesex* (*dsx*), and *fruitless* (*fru*) [2, 3]. An X:A ratio of 1 promotes transcription of *Sxl* and results in feminization, while an X:A ratio of 0.5 results in *Sxl* suppression and male differentiation [4, 5]. Sxl proteins control the splicing of female *tra* mRNA which gives rise to functional proteins, while no functional Sxl proteins are produced in the male [6]. Tra and tra2 proteins control the splicing of *dsx* and *fru* which are located at the bottom of the sex determination pathway to maintain sexual development and behavior [7–9]. The sex determining genetic cascade based on *tra*/*tra2* control of *dsx* and *fru* splicing is widely conserved in many Diptera, Coleoptera and Hymenoptera, while the domesticated *Bombyx mori*, lack *tra*/*tra2* as regulators of *dsx*/*fru* [3]. In the silkworm, the sex determination cascade involves at least 4 distinct components: a female-enriched PIWI-interacting RNA (*fem*), a responding gene, *BmMasc*, a P-element somatic inhibitor (*BmPSI*)/ (*BmImp*), and *Bmdsx* [10–12]. The product of the W chromosome derived *fem* piRNA targets the downstream gene, *BmMasc*, to control *Bmdsx* sex-specific splicing; *BmPSI* and *BmImp* regulate *Bmdsx* splicing through binding CE1 sequences of *Bmdsx* pre-mRNA [10]. Although the upstream signal is not conserved, with X:A and *fem* as the primary signals in the fruit fly and silkworm, respectively, the role of *dsx* is conserved.

Insects have sex-specific splicing that generates a male-(*dsx^M^*) and female-specific (*dsx^F^*) *dsx* isoform. Previous reports have shown that the *Bmdsx* gene products (*Bmdsx^M^* and *Bmdsx^F^*) control sexually dimorphic traits such as formation of abdominal segments and external and internal genitalia [13–16]. Studies of *D. melanogaster* have shown that the development of numerous sexually dimorphic traits are controlled by *Dmdsx*, including *Dmdsx^F^* controlled female-specific yolk gene transcription and female-specific spermathecae, *Dmdsx^M^* controlled male-specific abdominal pigmentation and male-specific sex combs [17–20]. The *Dmdsx^M^* gene products also regulate courtship behaviors, including licking, courtship song and copulatory behaviors [21]. The courtship behavior of *D. melanogaster* males consists of a series of discrete elements, including orientation toward the female, following the female, extending and vibrating one wing to produce a courtship song, licking the external genitalia, and attempting copulation [22]. *Dmdsx^M^* is expressed in about 700 neurons of the central nervous system, the majority of which also express *Dmfru^M^* [21]. *Dmfru^M^* function is both necessary and sufficient for nearly all aspects of male courtship behavior, and it is expressed in a dispersed subset of approximately 2,000 neurons in the central and peripheral nervous systems [23–25]. Males lacking *Dmfru^M^* appear to be normal externally yet are profoundly defective in most aspects of courtship behavior [22]. Moreover, *Dmdsx^M^* is necessary and sufficient for *Dmfru^M^*-independent courtship [26].

Although in recent years studies of neural and genetic mechanisms of sexual behavior in fruit flies indicate that *Dmdsx^M^* and *Dmfru^M^* are the master regulators of many sexually differentiated processes and behaviors [27, 28], how these master genes act to control neural development to build these complex behaviors by regulating downstream genes is still not fully understood [29]. Many studies have shown the *fru* gene to be conserved functionally with sex-specific splicing expression patterns among Diptera, including *Anopheles gambiae*, *Ceratitis capitata*, *Aedes aegypti*, *Nasonia vitripennis*, and *Musca domestica*, and a Blattodea, *Blatella germanica* [29–35]. However, the genetic regulatory mechanism of sexual behavior remains unclear in lepidopteran insects. Our previous study showed that loss of the *Bmdsx^M^* blocks male sexual behavior. The defective expression of *BmOR1* in male mutants of *Bmdsx^M^* contributes to the failed courtship behavior of orientation and leads to subsequent rejection of males by females [13].

*B. mori* has a simple sex pheromone system: female silkmoths emit the sex pheromones bombykol [(E,Z)-10,12-hexadecadien-1-ol] and bombykal [(E,Z)-10,12-hexadecadienal] at a typical ratio of 11:1. The major pheromone bombykol triggers the full sexual behavior of the male moth [36, 37]. Two sex pheromone receptors, BmOR1 and BmOR3, have male specific expression and are specific for bombykol and bombykal, respectively [38]. The pheromone binding protein BmPBP1, which has a male-biased expression, is required for the selectivity of BmOR1 for bombykol [39]. Loss of function mutations in *BmPBP1* or *BmOR1* cause the disappearance of male sexual behavior [40, 41]. Previous studies showed that the sex determination gene *Bmdsx* controls the expression of *BmPBP1* and of *BmOR1* [42, 14]. These results suggest that *Bmdsx* promotes male sex behavior, activating specific receptors of the olfactory system.

In the current study, we used genetic and electrophysiological approaches to investigate the molecular regulatory mechanisms controlling sexual behaviors in the silkworm by analyzing the mating behaviors of silk moths with mutations in known sex determination factors. Using our previously reported non sex-specific mutants *BmMasc*, *BmPSI*, and *Bmdsx* [12, 13], and the *Bmfru* mutant described here, we found that the sex determination pathway influences the development of the olfactory system to regulate courtship and mating behaviors. The sex determination pathway regulates morphological development and also the response to bombykol. The sexually dimorphic antennal morphologies of the male mutants of *BmMasc*, *BmPSI*, and *Bmdsx* are affected, which result in abnormal responses to bombykol and loss of courtship behavior. In contrast, male mutants of *Bmfru* show normal courtship but defective mating behavior. Knockout of the *Bmfru* downstream gene, *BmOR3*, impairs the response to bombykal but not bombykol, with normal courtship but extended mating behavior. Our data provide *in vivo* evidence of the function of the sexual differentiation pathway in the regulation of olfactory-based sexual behavior of *B. mori*.

## Results

### Sex determination pathway mutants have altered courtship and mating behaviors

We used a binary transgenic CRISPR/Cas9 system to obtain a *Bmfru* mutant. The *Bmfru* gene had not been reported in the silkworm, so we searched the silkworm genome database and identified a locus that encodes a protein (NCBI Accession number: EU649701.1) with a conserved BTB domain found in the FRU proteins of *D. melanogaster* and *M. domestica* (S1A Fig). We designed a small guide RNA (sgRNA) to target exon 3 of the putative *Bmfru* gene (S1B Fig). Through germline transformation, we obtained a single *U6-sgRNA* transgenic line. To obtain heritable and homozygous mutants to assess preference behaviors, we performed a series of crossing strategies and PCR-based screening experiments (S1C Fig) as described previously [43]. The *U6-sgRNA* and *nos-Cas9* parental transgenic lines were crossed with each other to obtain F_1_ founder moths, then ten random F_1_ female moths were crossed with wild-type males to obtain heterozygous mutants. F_2_ heterozygous mutant females were individually crossed with wild-type males to obtain independent lines of F_3_ heterozygous progeny, each potentially carrying a unique mutant allele. Individuals within F_3_ heterozygous lines were crossed with each other to obtain F_4_ homozygous mutants (S1C Fig) resulting in approximately 25% homozygous mutants in the F_4_ progeny. We characterized the mutations by PCR using gene-specific primer pairs. All of the homozygous mutants had four deleted base pairs that resulted in a premature termination codon (Fig S1D), confirming that the original CRISPR/Cas9-induced mutation had occurred.

Next, we analyzed courtship behavior of male mutants with a wild-type virgin female as the target and a mutant male (M-M) as the test at distances of 10 and 20 centimeters. The male sexual behavior of silk moth consists of the following steps: the male silk moth first recognizes the female by responding to a sex pheromone released by the female, the male moth then exhibits a programmed sequence of walking consisting of transitory bouts of straight-line walking, zig-zag turns and looping to climb toward the female. Then the male displays orientation, wing flapping (wing song), or turns around, and uses its forelegs to touch the female’s abdomen. After confirming the position of the female’s external gentitalia the male attempts to mate with the female using its claspers. Once the male mates with female, the male continues to flap its wings intermittently and copulates for several hours. The *Bmfru* mutant males did recognize wild-type females and displayed normal courtship behavior. Nevertheless, despite many attempts at copulating, the *Bmfru* mutant males could not mate with the wild-type female (S1-3 Movies). Thus, the courtship index of *Bmfru* mutant males was normal, whereas the mating index was zero (Fig 1). In contrast, *BmMasc*, *BmPSI*, and *Bmdsx* mutant males did not display any courtship behavior (i.e., no orientation, wing song, or turning around; *Bmdsx* behavior is illustrated in S4, S5 Movies), so the courtship and mating indexes of these three mutants were zero (Fig 1).

**Figure 1.**
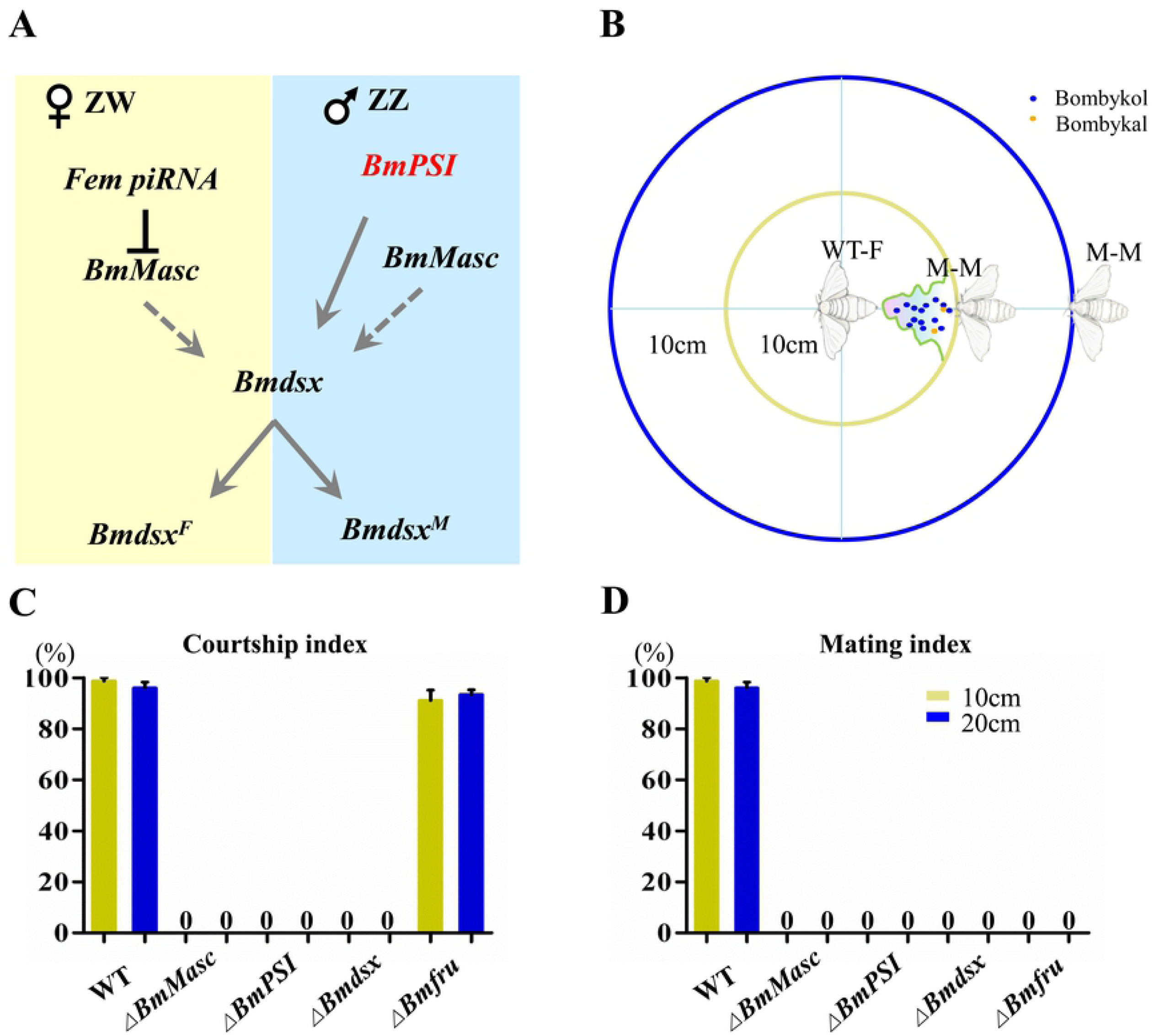
Loss of sex determination pathway genes impairs male courtship and mating behaviors. (A) Diagram of the sex determination cascade in the silkworm. (B) Diagram of the behavioral test setup. The adult male (M-M) is placed at a distance of 10 cm or 20 cm from a wild-type adult female (WT-F, which releases pheromones) or from a wild-type male control, and behavior is monitored. (C and D) The courtship and mating behavior indexes for wild-type males (WT) and *BmMasc*, *BmPSI*, *Bmdsx*, and *Bmfru* mutant males. The results are expressed as means ± SEM for 30 independent biological replicates each consisting of a single pair mating. An index of 0% indicates the absence of courtship behavior or failed mating behavior, whereas 100% indicates normal behavior.

### Antennal structures are abnormal in *BmMasc*, *BmPSI*, and *Bmdsx* mutants

Since insect antennae play a key role in the olfactory-based chemical communication necessary for courtship behavior, we evaluated the morphology of antennal structures in the mutants. In wild-type *B. mori*, the male antenna is larger and longer than the female antenna, and male antennae have greater numbers of sensilla trichoidea than female. The antennae were significantly shorter in males with mutations in *BmMasc* (18%), *BmPSI* (17%), and *Bmdsx* (9%), whereas antenna lengths were normal in *Bmfru* mutant males (Fig 2A and C). Using scanning electron microscopy (SEM), we observed that the numbers of sensilla trichoidea in a single field of view were significantly decreased in males with mutations in *BmMasc* (38%), *BmPSI* (40%), and *Bmdsx* (26%) compared to wild-type males, whereas the *Bmfru* male mutants had similar numbers of sensilla trichoidea to wild-type males (Fig 2B and D). The female mutant antennae had no significant change compared with wild-type females, except for *BmMasc-F* which had a slight increase in the length of antennae (Fig 2C). These results suggest that loss of *BmMasc*, *BmPSI*, or *Bmdsx* affects the development of antennal structures, which might underlie the observed dysfunctions in courtship and mating.

**Figure 2.**
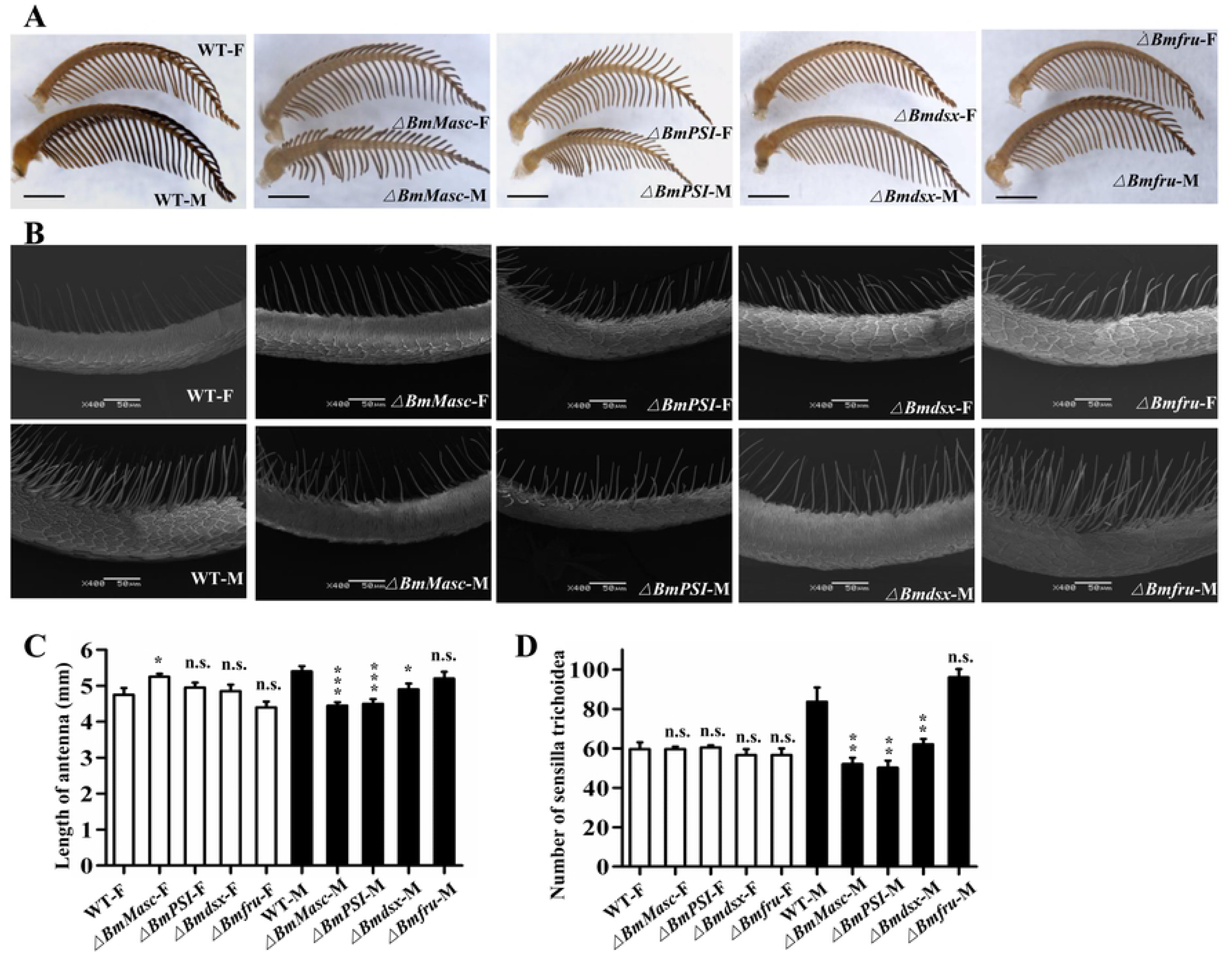
Silkworms with mutations in *BmMasc*, *BmPSI*, and *Bmdsx* have abnormal antennal structures. (A) Gross morphology of antennae of wild-type and mutant males (M) and females (F). Scale bars: 1 mm. (B) SEM images of the sensilla trichoidea structures in the middle of the antennae of wild-type and mutant males. Scale bars: 50 μm. (C) Antennal lengths of wild-type and mutant adults. The results are expressed as the means ± SEM of 10 independent biological replicates. * and *** represent significant differences at the 0.05 and 0.001 levels (ANOVAs), respectively, compared with the WT-F and WT-M; n.s. indicates that the difference is not statistically significant. (D) The number of sensilla trichoidea in one SEM scan field in wild-type and mutant adults. The results are expressed as the means ± SEM of 5 insects per group. ** represents significant differences at the 0.01 level (t-test) compared with the control; n.s. indicates that the difference is not statistically significant.

### Electrophysiological analyses reveal abnormal responses to sex pheromone components upon loss of sex determination genes

The female silkmoth attracts males by emitting sex pheromones. The pheromone bombykol is critical to attracting the male moths, whereas the minor pheromone bombykal plays an antagonistic role in mating behavior [36, 44]. We tested the responsiveness of mutant males to bombykol and bombykal using two methods: Electroantennogram (EAG) and single sensillum recording (SSR). We used EAG to detect the responsiveness to bombykol or bombykal at the level of whole antennae and SSR to evaluate responses of individual long sensilla trichoidea as described previously [40]. Compared to wild-type males and to *Bmdsx* and *Bmfru* mutants, the *BmMasc* and *BmPSI* mutants showed significantly lower EAG responses to bombykol (Fig 3A and C). *BmMasc*, *BmPSI*, and *Bmfru* mutant males displayed significantly lower EAG responses to the minor pheromone bombykal than *Bmdsx* mutants; nevertheless, the response of the *Bmdsx* mutants was also significantly lower than that of the wild-type moths (Fig 3A and D). These findings suggested that loss of any of the functions exerted by these sex determination and sexual differentiation genes interrupts the neuronal response to pheromones at the level of the antennae.

**Figure 3.**
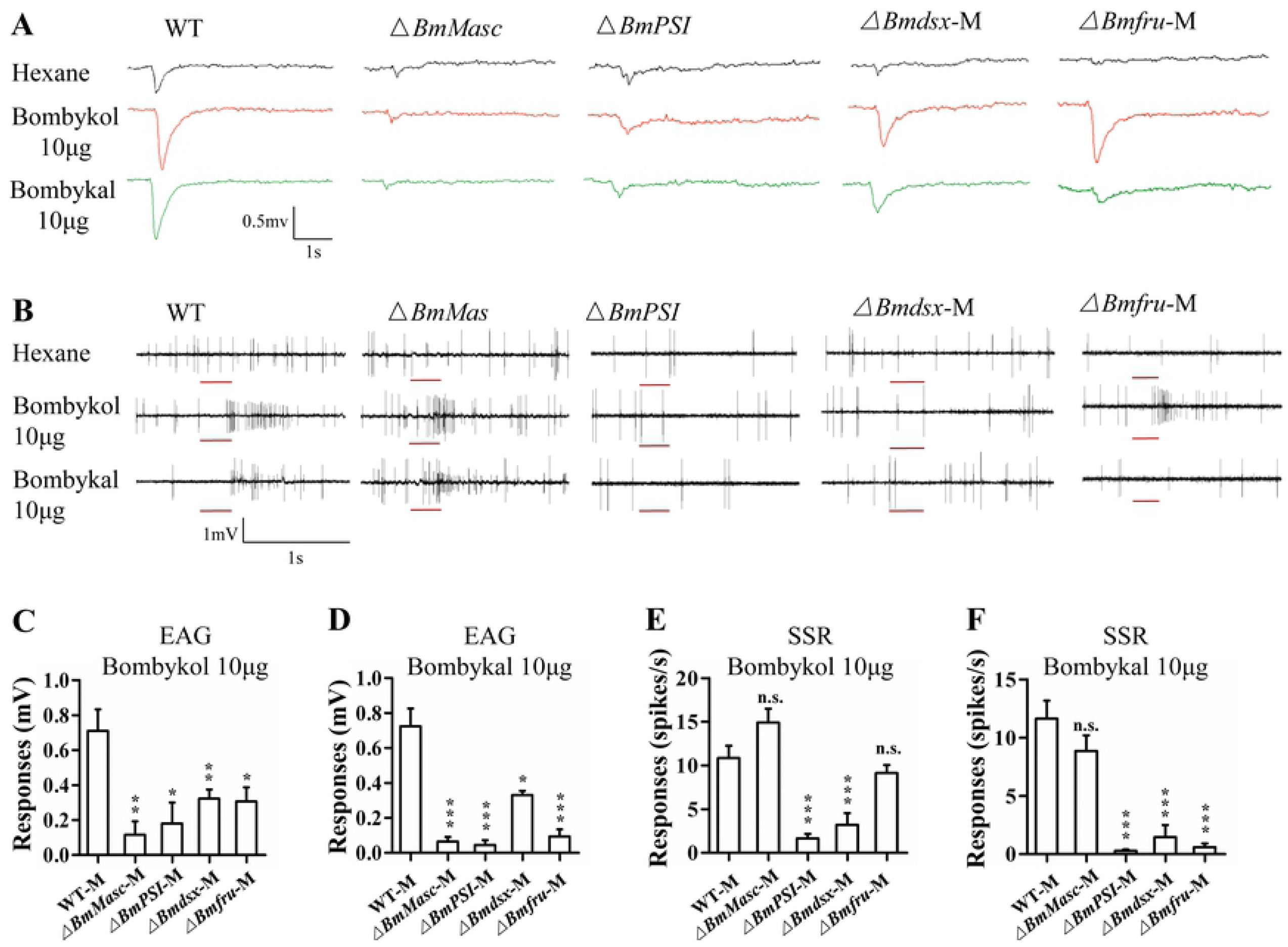
Electrophysiological analyses reveal abnormalities in responses to pheromone components of male silkworms with mutations in sex determination genes. (A) Representative EAGs of wild-type and mutant *BmMasc*, *BmPSI*, *Bmdsx*, and *Bmfru* male moths in response to hexane (upper panel), 10 μg bombykol (middle panel), and 10 μg bombykal (lower panel). (B) Representative single sensillum recording (SSR) from wild-type and mutant *BmMasc*, *BmPSI*, *Bmdsx*, and *Bmfru* males in response to hexane (upper panel), 10 μg bombykol (middle panel), and 10 μg bombykal (lower panel). The stimulus was applied for 300 ms, indicated by a red line under the trace. (C and D) Mean responses of male antennae to C) 10 μg of bombykol and D) 10 μg bombykal. The statistical significance between WT (n=10) and *BmMasc* (n=7), *BmPSI* (n=5), *Bmdsx* (n=8), and *Bmfru* (n=11) mutant responses was analyzed with ANOVAs. Data are shown as means ± SEM; *, **, and *** represent significant differences at the 0.05, 0.01, and 0.001 levels, respectively, compared with the WT-M as analyzed by an unpaired Student’s t-test. (E and F) Mean responses of neurons in male sensillum trichodea to E) 10 μg of bombykol and F) 10 μg bombykal. The statistical significance between WT (n=30) and *BmMasc* (n=50), *BmPSI* (n=44), *Bmdsx* (n=30), and *Bmfru* (n=76) mutants was analyzed with ANOVAs. Data are shown as means ± SEM; *** indicates p < 0.001, compared with the WT-M, and n.s. indicates no significance.

Next, we examined the responses of neurons within the long sensillum trichodea of the four mutants and wild-type males using SSR. From the spike traces of wild-type silkworms, two neurons (expressing *BmOR1* or *BmOR3*) were distinguished in the long sensillum trichodea. A larger amplitude was induced by the *BmOR1* neuron responding to bombykol, and a smaller one was evoked by the *BmOR3* neuron responding to bombykal (Fig 3B). The neuronal responses of the *BmPSI* and *Bmdsx* mutants to both bombykol and bombykal were significantly lower than wild-type whereas the neuronal responses of the *BmMasc* mutant to bombykol and bombykal were similar to wild-type males (Fig 3B, E and F). Although the single sensillum trichodea in the *BmMasc* mutants responded normally to bombykol and bombykal, it is possible that a decrease in the number of sensillum trichodea may contribute to the failure of courtship behavior of these mutants. The *Bmfru* mutants responded normally to bombykol but did not respond to bombykal (Fig 3B, E and F). This suggested that *Bmfru* mutants display normal courtship behavior because they have a normal response to bombykol.

### Expression of male-biased olfactory system genes is altered in mutants

Olfaction plays an important role in insect behaviors such as mate recognition. To test whether expression of olfactory system genes in the antennae were affected by loss of sex determination pathway genes, we quantified expression of genes that encode factors necessary for mate recognition by males, notably, pheromone binding proteins BmPBP1, BmPBP2, and BmPBP3 and odorant receptors BmOR1, BmOR2, and BmOR3 [45]. BmPBP1 binds to bombykol, and its dysfunction causes the failure of male courtship behavior [41]. Although *BmPBP1* was expressed at wild-type levels in males mutant in *Bmfru*, it was significantly decreased in the males mutant in *BmMasc*, *BmPSI*, and *Bmdsx*. Compared to wild-type males, *BmPBP2* and *BmPBP3* RNA expression levels were significantly higher in males mutant in *BmMasc*, *Bmdsx*, and *Bmfru* but lower in *BmPSI* mutants (Fig 4). Levels of *OR1* and *OR3* were significantly lower in all mutants (Fig 4). Mutations in *Bmdsx* reduce significantly the expression not only of *PBP1* and *BmOR1*, but also *BmOR3*, and increase the expression of *BmPBP2* and *BmPBP3*. No significant data are available for *BmOR2* in *Bmdsx* mutant. Furthermore mutations in *Bmfru* reduce significantly the expression of *BmOR1* and much more of *BmOR3*, and increase the expression of *BmPBP3*, and possibly also of *BmPBP2*. No significant data are available for *BmPBP1* and *BmOR2*. So the expression of *BmOR3* is increased in males by both *Bmdsx* and *Bmfru*, with *Bmfru* having an higher effect. *BmOR1* higher expression in males is promoted mainly by *Bmdsx* but also by *Bmfru* (Figure 4). *BmOR2* expression was normal except in the *BmPSI* mutant where it was expressed at levels lower than those observed in wild-type males (Figure 4). These expression patterns suggested that BmPSI is required in males for normal expression of 5 out of the 6 olfactory genes tested.

**Figure 4.**
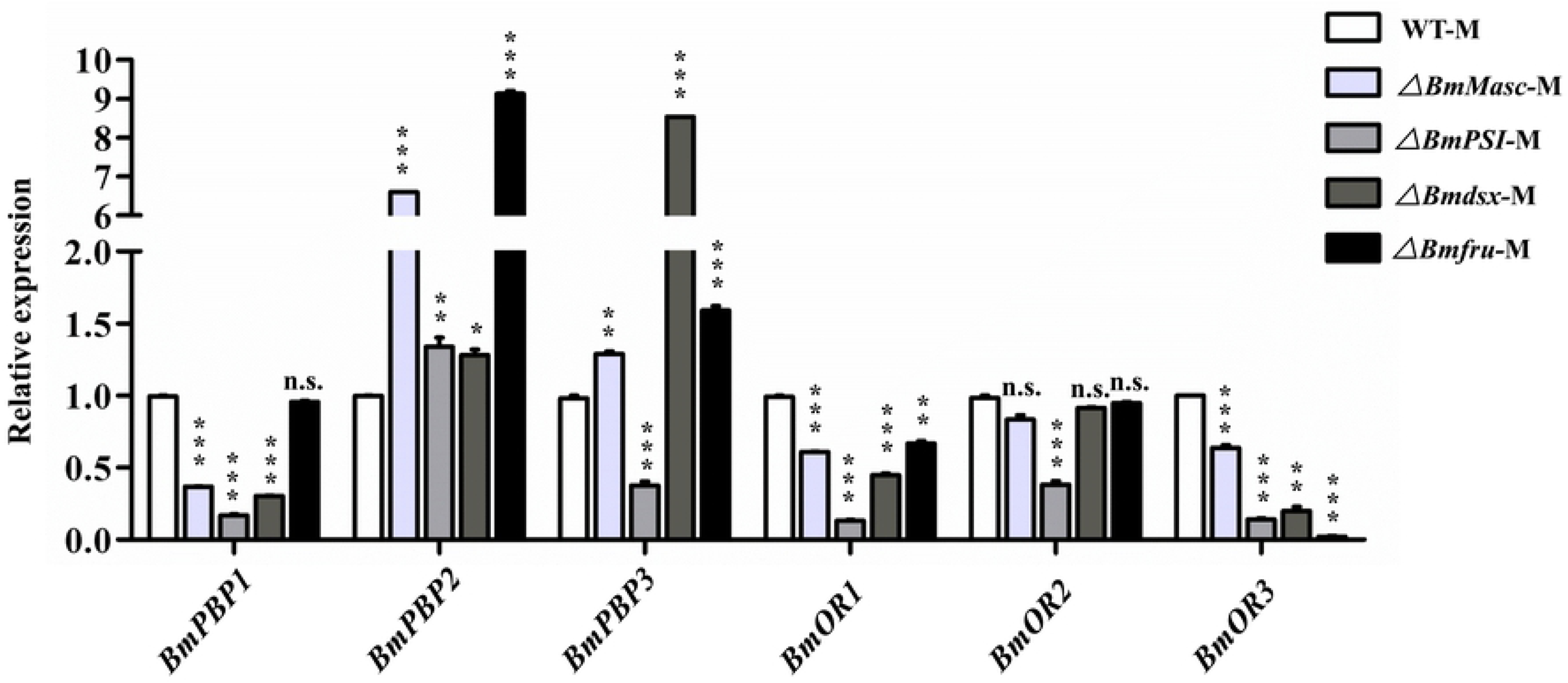
Olfactory sensory system genes are down-regulated in sex determination gene mutants. Relative mRNA expression levels of *BmPBP1*, *BmPBP2*, *BmPBP3*, *BmOR1*, *BmOR2*, and *BmOR3* in WT and mutant males. Three individual biological replicates were performed with real-time quantitative PCR (qPCR). Plotted are means ± SEM. ** and *** represent significant differences at the 0.01 and 0.001 levels (ANOVAs), comparing each gene was with the corresponding WT-M; n.s. represents not significant.

The normal response to sex pheromones by a single sensillum of the *BmMasc* mutant as shown by SSR may be due to the expression of *BmOR1* and *BmOR3* at about 60% of wild-type levels. Interestingly, expression of *BmOR3* was only 2% of wild-type levels in the *Bmfru* mutant, whereas the *BmOR1* gene was expressed at about 70% of levels in the wild-type males and *BmPBP1* expression was normal. Such relatively low expression of *BmOR1* and *BmPBP1* likely allowed the normal response to bombykol resulting in the normal courtship behavior of these mutant males. Altogether, these results suggest that sex determination pathway genes have an important role in establishing the sexually dimorphic expression of olfactory system genes, whereby *Bmdsx* primarily contributes to the expression of *BmPBP1* and *BmOR1* while *Bmfru* primarily contributes to the expression of *BmOR3*.

### Loss of *BmOR3* does not alter courtship behavior but does extend mating time

To further analyze the olfactory system-related genes in the antennae of *Bmfru* mutant males, we compared antennal transcriptomes of the adult *Bmfru* mutant male silk moths to those of wild-type males using RNA-seq. We identified 273 differentially expressed genes, 176 of which were down-regulated and 97 of which were up-regulated (S2A Fig). We found that olfactory system-related genes including *BmPBP1*, *BmPBP2*, *BmPBP3*, *BmOR1*, and *BmOR3* were differentially expressed (S2B Fig) as shown by RT-qPCR. In the *Bmfru* mutant antennae, *BmOR1* expression was 0.99-fold lower and *BmOR3* was 5.32-fold lower than in the wild-type antennae.

To further investigate the function of *BmOR3*, we used a binary transgenic CRISPR/Cas9 system to target exon 2 of *BmOR3* (S3A Fig). We established three independent U6-sgRNA parental transgenic lines and then made crosses within each line to obtain F_1_ founder moths. Quantitative real-time RT-PCR showed a decreased in *BmOR3* mRNA levels of F_1_ founder individuals by over 99% in each of the three mutant lines compared with wild type moths (S3B Fig). Characterization of the somatic mutations by PCR using gene-specific primer pairs indicated that mutants had deletions at the target site caused by non-homologous end joining-induced indels (S3C and D Fig). These results suggested that F_1_ individuals carried truncated proteins of BmOR3, so we used the F_1_ founder moths to analyze adult behavior.

The *BmOR3* mutant males displayed normal courtship behavior, including orientation, wing song, and turning, and the courtship and mating indexes were only slightly below normal (Fig 5A and B). As noted previously, the normal duration of copulation is several hours. Although almost all wild-type males had autosegregated after 12 hours of mating, most of the *BmOR3* mutant males had not (Fig 5C). Electrophysiological analyses revealed that the *BmOR3* mutants responded normally to bombykol but had lost responsiveness to bombykal (Fig 6); this was also the case for the *Bmfru* mutants. These findings indicated that *BmOR3* is not necessary for courtship.

**Figure 5.**
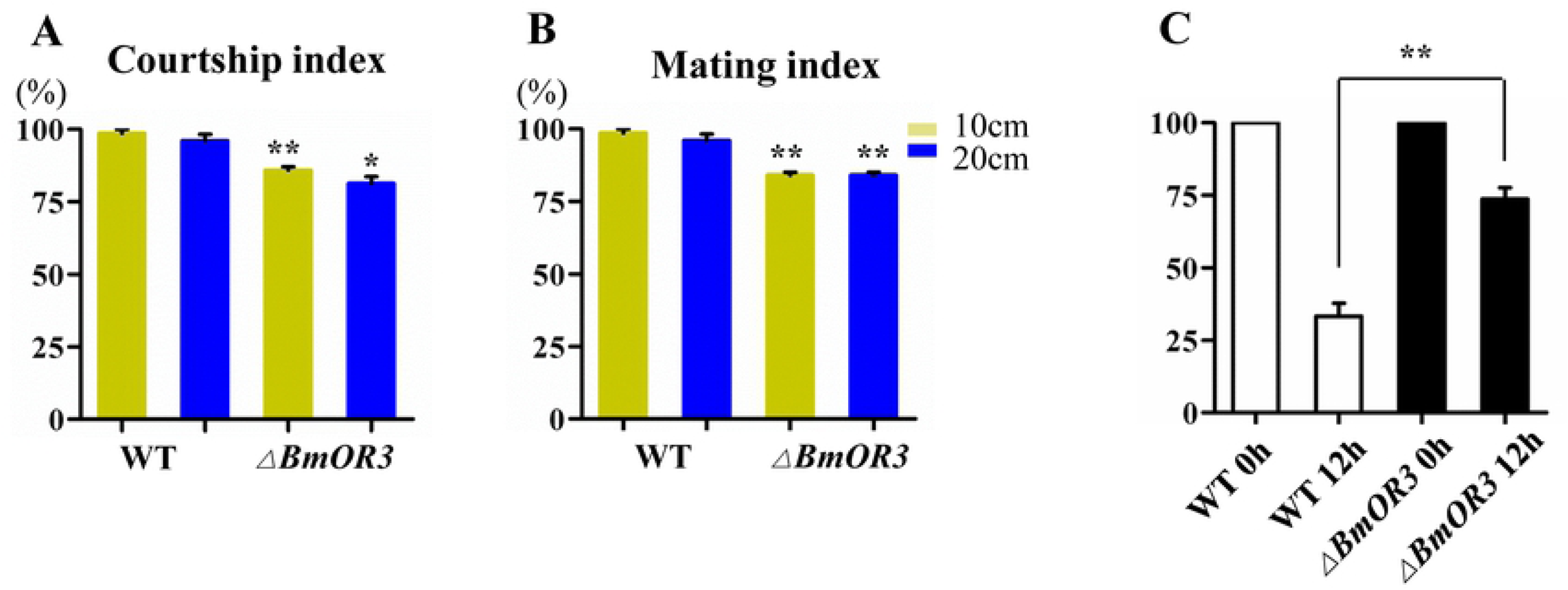
Loss of *BmOR3* expression extends mating time. (A and B) Courtship and mating behavior indexes of *BmOR3* mutant and WT males. Data are shown as mean ± SEM of 50 independent biological replicates. (C) Percentage of autosegregated WT and mutant males at 0 h and 12 h. ** indicates significant difference at the 0.01 level.

**Figure 6.**
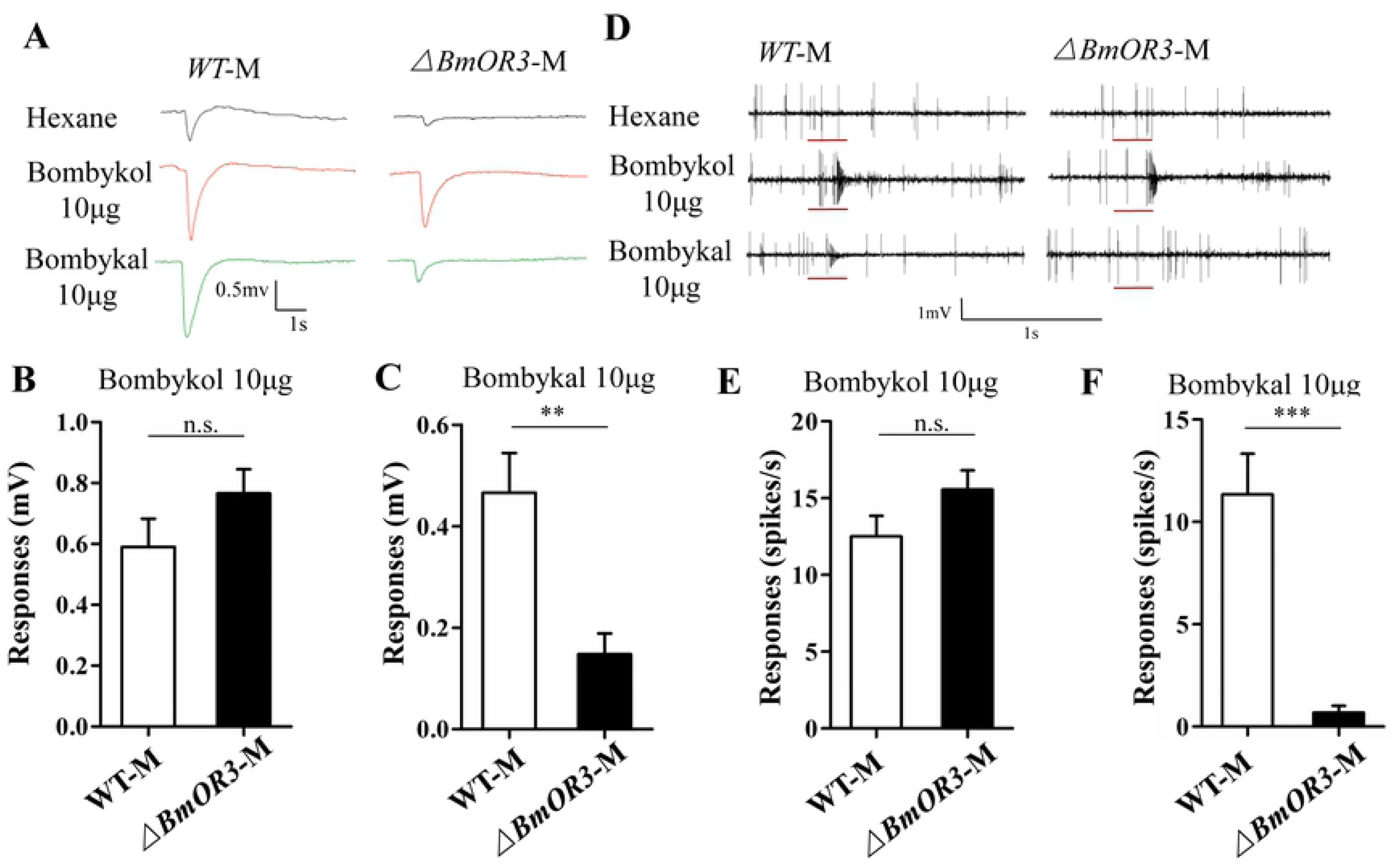
*BmOR3* mutant male silk moths have nearly normal electrophysiological responses to bombykol but not bombykal. (A) Representative EAGs from wild-type and *BmOR3* mutant male moths in response to hexane (upper panel), 10 μg bombykol (middle panel), and 10 μg bombykal (lower panel). (B and C) Mean responses of WT (n=9) and *BmOR3* mutant (n=9) male antennae to B) 10 μg of bombykol and C) 10 μg of bombykal. Data are means ± SEM; n.s. indicates no significant difference and ** represents a significant difference at the 0.01 level as determined by Student’s t-test. (D) Representative single sensillum recording (SSR) of wild-type and *BmOR3* mutant male moths in response to hexane (upper panel), 10 μg bombykol (middle panel), and 10 μg bombykal (lower panel). The stimulus was applied for 300 ms as indicated with a red line under the trace. (E and F) Mean (± SEM) responses of neurons in male sensillum trichodea to E) 10 μg of bombykol or F) 10 μg of bombykal in WT (n=26) and *BmOR3* mutant (n=40) male moths. *** indicates p < 0.001 and n.s. indicates no significant difference as determined by Student’s t-test.

## Discussion

Previous studies have shown that *BmOR1* and *BmOR3* encode sex pheromone receptors in the silkworm [37, 39, 40]. Although the mechanism of sex determination in the silkworm has also been studied in recent years [13, 46, 47], it has not revealed a connection between these two major pathways which control the sexual behavior and morphology of the insect. Here, we provide genetic evidence for a functional interplay between these sex pheromone receptors and genes in the sex determination pathway. Our results support the notion that the *Bmdsx*-*BmOR1*-bombykol pathway regulates courtship behavior and that *Bmfru*-*BmOR3*-bombykal regulates mating (Fig 7).

**Figure 7.**
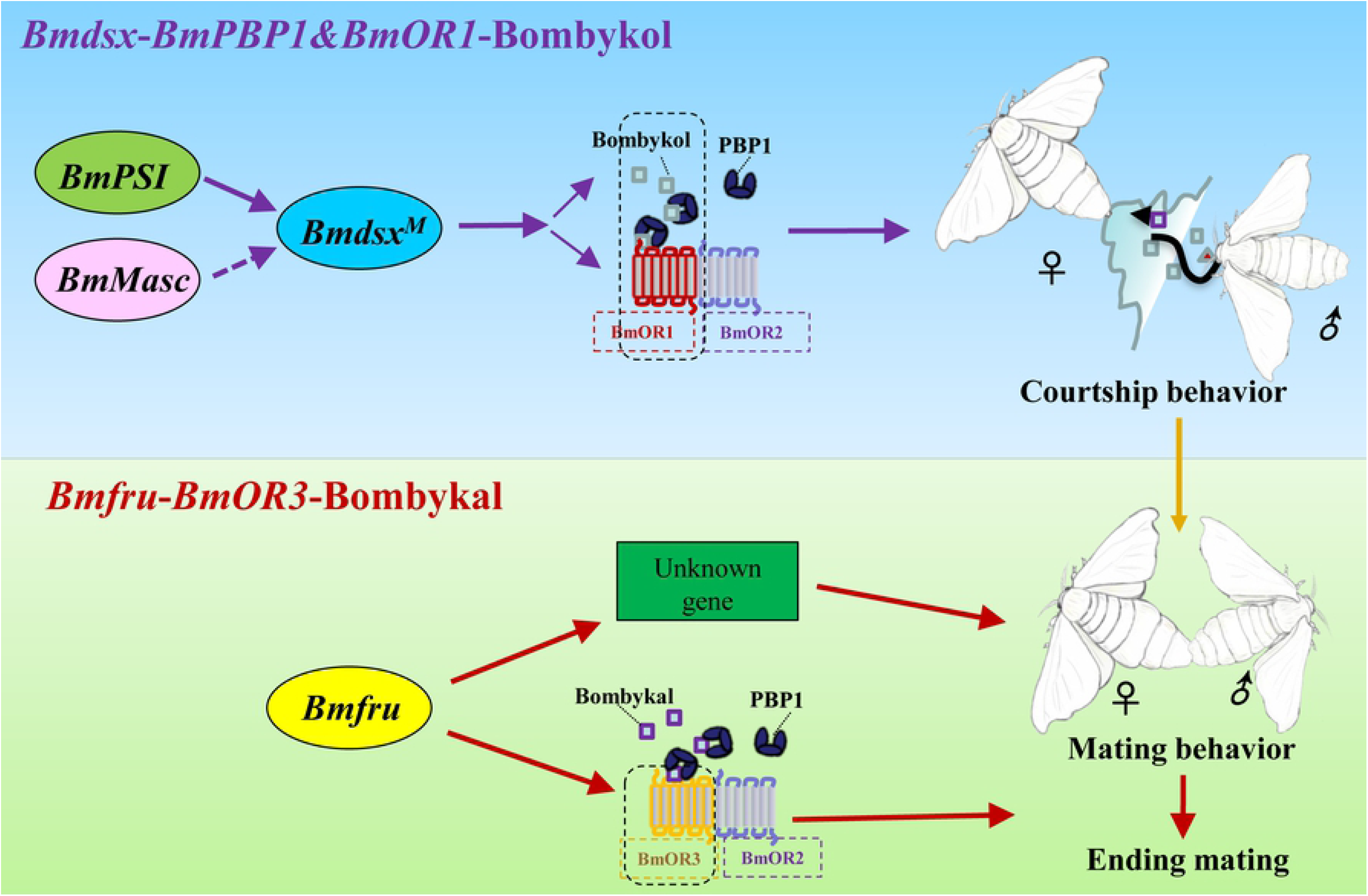
Proposed genetic regulation pathway of sexual behavior in *B. mori*. Sex determination pathway factors, olfactory sensory factors, and sex pheromones influence courtship and mating behavior. The Bmdsx-BmOR1/3-Bombykol and Bmfru-BmOR3-Bombykal cascades are the two primary pathways involved in olfactory-based sexual behavior. Disruption of the sex pathway gene *Bmdsx* blocks the expression of both *BmOR1* and *BmOR3*, whereas disruption of *Bmfru* mainly inhibits expression of *BmOR3*. Thus, mutation of *Bmdsx* leads to an abnormal response to bombykol and bombykal and inhibits courtship and mating by male moths, but mutation of *Bmfru* disrupts termination of mating due to an abnormal response to bombykal.

Exposure to bombykol, the major female pheromone, was sufficient to induce pheromone source orientation behavior in male moths, as was the artificial activation of *BmOR1* expressing olfactory receptor neurons [48, 49]. Previous studies have shown that *BmOR1* and *BmOR3* are specifically expressed in adult males [50]. This sexually dimorphic pattern of gene expression suggested that *BmOR1* and *BmOR3* are regulated by *Bmdsx* and *Bmfru*. TALEN-mediated knock out of *BmOR1* indicated that BmOR1 is the sex pheromone receptor that mediates the pheromone response in male silkmoths [40]. The minor component, bombykal, was thought to negatively modulate the initiation of orientation behavior [44]. However, our results support a conclusion that BmOR3 has little effect on the initiation of orientation behavior. In our study, *BmOR3* mutants which did not respond to bombykal had normal courtship but had extended mating behavior. Additionally, *Bmfru* mutants, which expressed almost no *BmOR3*, had normal courtship behavior but did not mate. This suggests that the *BmOR3*-bombykal interaction does not play a key role in the recognition of females by male moths and that *Bmfru* might act through other pathways to control mating behavior. Daimon *et al.* showed that the wing fluttering response of *B. mori* males to bombykol is strongly inhibited by bombykal, thereby indicating that bombykal acts as a behavioral antagonist [44]. All of these observations suggest that bombykal may play role in terminating copulation behavior.

In dipteran insects, *fru* is located downstream of *tra* in the sex determination pathway, and *dsx* is involved in the regulation of sexual behavior [34, 51]. No gene homologous to *tra* has been found in the silkworm genome, indicating that the sex determination pathway in *B. mori* is different from dipterans. Additionally, *Bmfru* is not like *Dmfru*, which has sex-specific splicing. *Bmfru* was found to have male-baised expression in the brain and testis [52]. This suggests that the regulatory mechanisms involving *fru* gene expression are different between fruitfly and silkworm. In *Drosophila*, a PSI–U1 snRNP interaction regulates male mating behavior, and one of its direct targets is *Dmfru* [53]. This suggests that *DmPSI* regulates mating behavior through *Dmfru* and other genes. By contrast, in the silkworm, *BmPSI* and *BmMasc* are upstream of the sex determination factor *Bmdsx* [11, 13]. Knockout of *BmPSI* caused the failure of courtship. In addition to *BmOR2*, the genes encoding *BmOR1*, *BmOR3*, *BmPBP1* and *BmPBP*, were downregulated in the *BmPSI* mutant males produced in our study. This suggested that BmPSI might exert pleiotropic effects on olfactory development. On the other hand, *BmOR2* was expressed normally in *BmMasc* mutant males, and changes in expression of other olfactory system genes were more moderate in *BmMasc* mutant males than in *BmPSI* mutant males. Moreover, the *BmMasc* mutant responded to pheromones normally as shown by SSR, even though the signal from whole antennae was decreased as shown using EAG. These results suggest that olfactory neurons are functionally normal but may be decreased in number in the *BmMasc* mutant.

In conclusion, by using a comprehensive set of knockout mutations in genes *Bmdsx*, *BmPSI*, *BmMasc*, *Bmfru*, and *BmOR3*, we showed that the sex determination cascade in *B. mori* contributes to the establishment of olfactory system-based sexual behaviors. We found that the sex determination genes control behavior by regulating expression of genes encoding olfactory receptors. *BmMasc* and *BmPSI* act as upstream signals and *Bmdsx* acts as the performer located at the bottom of the cascade to control courtship behaviors by regulating *BmOR1*, as well as *BmOR2* (*BmPSI*) and both copies of *BmOR3*. The *Bmfru* gene product contributes to courtship behavior by regulating *BmOR3*, *BmOR1* (slight effect) and other unknown pathways. Future work should focus on how these sex determination genes affect neuronal modulations that influence sexual behaviors.

## Materials and methods

### Silkworm strain

Silkworms of the Nistari genetic background (a multivoltine, non-diapausing strain) were used in all experiments. Larvae were reared on fresh mulberry leaves under standard conditions. *BmMasc*, *BmPSI* and *Bmdsx* mutants are described in our previous reports. The parental transgenic U6-sgRNA and *nos-Cas9* lines were reared separately. Crossing these two lines produce heteroallelic mutations in somatic or germ cells of F_1_ individuals. F_1_ individuals carrying heteroallelic mutations were used in this study. The detection of genomic mutations and measurement of mRNA levels among these three F_1_ lines were reported previously [12, 13].

### Plasmid construction and germline transformation

To target the *Bmfru* and *BmOR3* genes, plasmids pBac[IE1-DsRed2-U6-sgRNAs] (*U6-sgRNA*) were constructed to express sgRNA under the control of the silkworm *U6* promoter and the DsRed fluorescence marker gene, under control of an *IE1* promoter. The sgRNA targeting sequences were selected by manually searching genomic regions for sequences that matched the 5′-GG-N_18_-NGG-3′ rule [54]. sgRNA sequences were checked bioinformatically for potential off-target binding to the relevant silkworm genomic sequence using CRISPRdirect (http://crispr.dbcls.jp/) [55]. All sgRNA and oligonucleotide primer sequences for plasmid construction are listed in Table S1. Plasmid construction was performed as described previously [13]. Each *U6-sgRNA* plasmid was mixed with a *piggyBac* helper plasmid and microinjected separately into fertilized eggs at the pre-blastoderm stage. G_0_ adults were mated to WT moths, and the resulting G_1_ progeny were scored for the presence of the DsRed marker gene product using fluorescence microscopy (Nikon AZ100).

### Genotyping analysis

Animals from each U6-sgRNA transgenic line (one *Bmfru* line and three *BmOR3* lines) were mated with the *nos-Cas9* line to obtain mutated F_1_ animals. Genomic DNA of mutated animals was extracted at the larval stage using standard SDS lysis-phenol treatment after incubation with proteinase K, followed by RNase treatment and ethanol precipitation. The resulting individual DNA samples from mutant animals were separated by sex using PCR amplification with primers specific to the W chromosome (S1 Table). Mutation events were detected by PCR amplification using gene-specific primers that bound upstream or downstream from each target (S1 Table). Amplified products were visualized by agarose gel electrophoresis. Amplicons were sub-cloned into the pJET-1.2 vector (Fermentas), and six positive clones of each line (one *Bmfru* and three *BmOR3* lines) were selected and sequenced using Illumina NextSeq 500 platform (Sunnybio, Shanghai).

### Photography and scanning electron microscopy (SEM)

Antennae of mutant and wild-type animals were dissected, photographed under a light microscope (Nikon, Tokyo, Japan) using a digital camera (Nikon DS-Ri1, Japan), and lengths measured on the images..

The dissected antennae were fixed overnight in a solution of 90 ml 70% ethanol, 5 ml acetic acid, and 5 ml 37% methyl aldehyde, dehydrated in a series of 70%, 80%, 90%, and 100% ethanol baths for 5 min each, and dried (CO_2_ for 6 h in a Critical Point Dryer). The dissected materials were coated with gold by JFC-1600 sputter (JEOL, Rigaku, Japan), and the middle parts of the antennae were observed by SEM using a JSM-6360LV microscope (JEOL, Rigaku, Japan).

### Analysis of courtship and mating behavior

*BmMasc*, *BmPSI*, *Bmdsx*, and *BmOR3* mutant males were from heterozygous F_1_ individuals. *Bmfru* mutant males could be distinguished by a behavior test monitoring in the presence of WT virgin females. In the absence of being able to fly, silkworm sexual behavior is dependent on walking distance. Therefore, we set a test field to measure movement within a radius of 10 cm or 20 cm as reported in a previous study [43] which was sufficient for male silk moths to recognize females. Once the male mates with the female, it continues to flap its wings and can remain copulated for several hours. Experientially, continuous mating for 30 minutes ensures normal sperm transfer and reproduction. So we defined and evaluated a courtship index by the following steps: (1) male moves toward the female and successfully displays orientation behaviors; (2) wing song; and (3) reorientation and tipping the abdomen. Courtship index was recorded as 1 when male moth displayed these three steps. The mating index was evaluated by measuring whether the male copulated with the female continuously for 30 minutes. The behavioral assays were performed at 25 °C and 60% relative humidity under normal ambient light.

### Electroantennogram (EAG) recordings

Five to eleven antennas each taken from more than five virgin male moths 1-2 days after eclosion were used for EAG recordings. EAG values were recorded by using a method similar to one previously reported [56]. The antenna was cut off at the base from the head, and a few terminal segments of the antenna were excised to achieve better contact. The cut ends of the antenna were connected with two recording glass electrodes filled with 0.1 M KCl. Five to eleven individuals of each genotype were evaluated. The recordings were performed under an Olympus SZ61 microscope.

Pheromone components (bombykol and bombykal, 96% purity, purchased from Nimrod Inc., Changzhou, China) were diluted in hexane (98% purity, Sigma-Aldrich Co., St. Louis, MO, USA) and a dose of 10 µg was used for each trial as previously described [43]. Hexane was used as the control. Briefly, a filter paper strip (2.5 × 0.9 cm) was wet with 10 μl test solution and allowed to dry for 3 min, then the paper strip was inserted into a Pasteur pipette placed perpendicularly through a hole in a metal-lined tube with a humid airflow of 0.5 L/min.

Signals were amplified 10-fold (10 s, starting 1s before stimulation) by a high impedance pre-amplifier (IDAC-2 USB System, Syntech, Kirchzarten, Germany) then sent to a computer via an analog-digital converter. Off-line analysis was carried out by EAGpro 2.0 software (Syntech, The Netherlands). Relative EAG responses for each compound were calculated by subtracting EAG response for the blank from the EAG response to the test compound. Data are presented as the means ± standard error of the mean (SEM) and were compared using an unpaired Student’s t-test.

### Single sensillum recordings (SSR)

To perform single sensillum recordings, a 1-2 day-old virgin male silk moth was placed in a remodeled 1-ml plastic pipette with the protruding head fixed by dental wax. The exposed antenna was attached to a cover-slip with double-faced adhesive tape. A recording tungsten wire electrode was inserted into the sensilla and reference tungsten wire electrodes were inserted into the compound eyes. Data were obtained for 3-6 individuals for each genotype and 10-20 sensilla for each individual were examined. The recordings were performed under a LEICA Z16 APO microscope at 920 × magnification.

Pheromone components were prepared as described in the “Electroantennogram (EAG) recordings” section. Humid air flow was set at 1.4 L/min, and a stimulus air pulse for 300 ms was controlled by a Syntech Stimulus controller (CS-55, Syntech, Kirchzarten, Germany). Signals were amplified 10-fold (10 s, starting 1 s before stimulation) by a high impedance pre-amplifier (IDAC-4 USB System, Syntech, Kirchzarten, Germany) then output to a computer via an analog-digital converter. Off-line analysis was carried out using AUTOSPIKE, v. 3.9, software (Syntech, Kirchzarten, Germany). The filter setting was 300 Hz at low cutoff and 2 kHz at high cutoff. The responses were measured by counting the number of action potentials within 1 s after stimulation. The number of olfactory sensory neurons housed in a single sensillum was determined based on the differences in spike amplitudes. Data are presented as the means ± SEM and compared with an unpaired Student’s t-test.

### Quantitative RT-PCR

Total RNA was extracted from 10 silkworm antennae from 5 males per genotype using Trizol reagent (Invitrogen) and treated with RNase-free DNAse I (Ambion). cDNAs were synthesized using the Omniscript Reverse transcriptase kit (Qiagen) in a 20 µl reaction mixture containing 1 µg total RNA. Quantitative real-time RT-PCR (RT-qPCR) assays were performed using SYBR Green Realtime PCR Master Mix (Thermo Fisher Scientific) on an Eppendorf Real-time PCR System MasterCycler RealPlex instrument. RT-qPCR reactions were carried out with gene-specific primers (Table S1). A 10-fold serial dilution of pooled cDNA was used as the template for standard curves. Quantitative mRNA measurements were performed in three independent biological replicates, and data were normalized to the amount of *Bmrp49* mRNA [12].

### RNA-seq protocol and data analysis

Illumina sequencing as perfomered as our previous study [57], total RNA was isolated from 10 *Bmfru* mutant and WT antennas using TRIzol (Invitrogen, Carlsbad, CA, USA), and the residual DNA was removed with RNase-free DNase I (New England BioLabs, Ipswich, MA, USA) for 30 min at 37 °C. For RNA-seq, library construction and sequencing using an Illumina HiSeq 2000 were conducted by BGI Genomic Services (Shenzhen, China), briefly described as follows. The mRNA was enriched using oligo (dT) magnetic beads samples were mixed with a fragmentation buffer, and the mRNA reduced to short fragments (∼200 bp). The first strand of the cDNA was synthesized using random hexamer primers, buffer, dNTPs, RNase H, and DNA polymerase I were added to synthesize the second strand, and the double-stranded cDNA was purified with magnetic beads followed by performing end repair and 3’-end single nucleotide adenine addition. Finally, sequencing adaptors were ligated to the fragments which were enriched by PCR amplification, and an Agilent 2100 Bioanaylzer and an ABI Step One Plus Real-Time PCR System were used to quantify libraries. The library products were sequenced using an Illumina HiSeq 2000 (BGI Biotech Co. Ltd.). The raw sequencing data were qualified, filtered, and mapped to the reference silkworm genome database (http://silkworm.genomics.org.cn/) using tophat/bowtie2. The UniGene abundances were measured in fragments per kb of exon per million fragments mapped (FPKM). The differentially expressed genes were annotated functionally using Gene Ontology and Kyoto Encyclopedia of Genes and Genomes annotations.

### Statistical analysis

Behavioral and RT-qPCR data were analyzed in GraphPad Prism 5, and electrophysiological data were analyzed in Spike2 and SigmaPlot. Experimental data were analyzed with the ANOVAs or Student’s t-test. At least three independent replicates were used for each treatment and means ± SEM were plotted. Detailed statistical information relating to each experiment is provided in the relevant Method Details or figure legends.

## Supporting Information

**S1 Fig. A binary transgenic CRISPR/Cas9 system induces homozygous mutations at the *Bmfru* locus in *B. mori*.** (A) The BmFRU protein, which contains the BTB domain conserved in dipteran insects, *D. melanogaster* and *M. domestica*. (B) Schematic representation of the exon/intron boundaries of the *Bmfru* gene. Exons are shown as boxes. Untranslated regions are shown as black boxes and coding regions as open boxes. Thin lines represent the introns and numbers are the lengths in kilobase pairs (kb). Target site locations are noted and PAM sequences are shown in red. (C) Crossing scheme to produce homozygous mutations. The binary transgenic CRISPR/Cas9 system in this study contains two lines, one of which contains the full *Cas9* ORF driven by the *nanos* (*nos*) promoter, and the other contains a U6 promoter-driven sgRNA. These two lines also encode the reporter genes *EGFP* and *DsRed2*, respectively. The two transgenic lines were crossed to produce founder animals that express both Cas9 and *Bmfru* sgRNAs. The founder female silkworms were backcrossed with wild-types to obtain heterozygous offspring (F_2_, *Fru*^+/-^). F_2_ heterozygous mutant females were individually crossed with wild-type males to obtain distinct F_3_ heterozygous lines. F_3_ moths heterozygous for the mutations were sib-mated to generate independent lines of homozygous animals (F_4_, *Fru*^-/-^). (D) Homozygous mutations confirmed by sequence analysis. The targeting sequence is shown in blue and the PAM sequence in red. The deleted base pairs (bp), ATGC, are indicated by the broken line.

**S2 Fig. Comparison of *Bmfru* mutant and wild-type male moth antennae transcriptomes.** (A) Plot of significantly differentially expressed genes in 10 mixed *Bmfru* mutant male antennas compared to 10 mixed WT adult male antennas. False discovery rate (FDR) was used to determine the threshold of p values in multiple tests. We use FDR ≤ 0.001 and the absolute value of log_2_Ratio ≥ 1 as thresholds to determine significant differences in gene expression. Yellow represents up-regulated genes, blue represents down-regulated genes, and gray represents genes without significant differences. (B) Olfactory sensory system genes with changes significant at p<0.05.

**S3 Fig. A binary transgenic CRISPR/Cas9 system induces loss of functional mutations at the *BmOR3* locus.** (A) Structure of the *BmOR3* gene with nine exons indicated by boxes (black boxes, 5’-and 3’-UTRs; white boxes, coding exons). Target sites 1 and 2 are binding sites for sgRNAs. (B) The mean transcript levels (± SEM) of *BmOR3* are down-regulated significantly compared to wild-type levels in the three *BmOR3* mutant male (M) and female (F) lines. At least five males with mixed antenna were examined for each line. *** indicates p < 0.001 compared with the relevant control using Student’s t-test. (C) Somatic mutations were induced in the F_1_ founder animals following crosses of nos-Cas9 with U6-sgRNA strains. PCR analyses with primers to amplify a region of 600 bp revealed deletion mutations in the G_0_ mutants. The red arrowhead indicates the deleted region. (D) Deletion mutation in the heterozygous offspring after crossing nos-Cas9 and U6-BmOR3sgRNA transgenic silkworm lines. The targeting sequence is shown in black, and the PAM sequence is in red. The deletion size in nucleotides is indicated above the red arrow at the site of the deletion.

**S1 Movie. A wild-type male successfully copulating with a wild-type female.**

**S2 Movie. A *Bmfru* mutant male failing to copulate with a wild-type female.**

**S3 Movie. A *Bmfru* mutant male failing to copulate with a *Bmfru* mutant female.**

**S4 Movie. A *Bmdsx* mutant male failing to copulate with a wild-type female.**

**S5 Movie. A *Bmdsx* mutant male failing to copulate with a *Bmdsx* mutant female.**

**S1 File. Protein sequences used in Figure S1.**

**S1 Table. Oligonucleotide primers used in this study.**

**S2 Table. Raw data for plots in** Fig’s 2 and 5.

**S3 Table. Raw data for plots in** Fig 2. The workbook contains two sheets. Sheet 1 shows data for antennal length. Sheet 2 presents data for the number of sensilla trichoidea in a single SEM scan field.

**S4 Table. Raw data for plots in** Fig’s 3 **and ig 6.** The workbook contains six sheets. Each sheet contains raw data for a separateEAG experiment.

**S5 Table. Raw data for plots in** Fig’s 3 **and 6.** The workbook contains six sheets. Each sheet contains raw data for a separate SSR experiment.

**S6 Table. Raw data for plots in** Fig 4. Q-PCR data showing the mRNA level for different mutations compared to WT.

**S7 Table. Raw data for plots in S3 Fig.** Q-PCR data showing level of *OR3* mRNA relative to *RP49*.

## Acknowledgments

We thank Jiqin Li and Xiaoyan Gao for SEM technique support. We thank all members of the Huang lab and Prof. Anjiang Tan for technical assistance and helpful discussions. We would like to thank Dr. Jacqueline Wyatt and Prof. Marian Goldsmith for proof reading the manuscript. We thank three anonymous reviewers for their constructive comments.

## Data Availability

The transcriptome data are available from the Dryad database (DOI: https://doi.org/10.5061/dryad.3xsj3txbw).

